# ESCRT-I controls lysosomal membrane protein homeostasis and restricts MCOLN1-dependent TFEB/TFE3 signaling

**DOI:** 10.1101/2021.07.26.453823

**Authors:** Marta Wróbel, Ewelina Szymańska, Noga Budick-Harmelin, Krzysztof Kolmus, Krzysztof Goryca, Michalina Dąbrowska, Agnieszka Paziewska, Michał Mikula, Jarosław Cendrowski, Marta Miączyńska

## Abstract

Within the endolysosomal pathway in mammalian cells, ESCRT complexes facilitate degradation of proteins residing in endosomal membranes. Recent studies revealed that yeast ESCRT machinery also sorts ubiquitinated proteins from the vacuolar membrane for degradation in the vacuole lumen. However, whether mammalian ESCRTs perform a similar function at lysosomes remained unknown. Here, we show that ESCRT-I restricts the size of lysosomes and promotes degradation of proteins from lysosomal membranes, including MCOLN1, a Ca^2+^ channel protein. Upon ESCRT-I depletion, the lysosomal accumulation of non-degraded proteins coincided with elevated expression of genes annotated to cholesterol biosynthesis and biogenesis of lysosomes, indicative of response to lysosomal stress. Accordingly, the lack of ESCRT-I promoted abnormal cholesterol accumulation in lysosomes and activated TFEB/TFE3 transcription factors. Finally, we discovered that in contrast to basal TFEB/TFE3 signaling that depended on the availability of exogenous lipids, the stress-induced activation of this pathway was Ca^2+^-MCOLN1-dependent. Hence, we provide evidence that ESCRT-I is crucial for maintaining lysosomal homeostasis and we elucidate mechanisms distinguishing basal from lysosomal stress-induced TFEB/TFE3 signaling.

## Introduction

Lysosomes are acidic organelles of animal cells that serve as a major degradative compartment for proteins or lipids delivered by endocytosis or autophagy (Luzio *et al*, 2007), hence are equivalent of a yeast vacuole. Besides providing cells with metabolites derived from degradation and with molecules taken up by endocytosis, lysosomes act also as signaling organelles (Ballabio, 2016). Reduced cargo delivery to lysosomes or their dysfunction causing lysosomal stress induce lysosome-related signaling pathways that adjust cellular metabolism (Ballabio, 2016). These pathways are orchestrated by kinases activated from the lysosomal surface or by changes in efflux of metabolites or ions from the lysosomal lumen. For instance, the release of calcium ions (Ca^2+^) from lysosomes via channels formed by mucolipin 1 protein (MCOLN1, also known as TRPML1) activates calcineurin, calcium-dependent phosphatase (Medina *et al*, 2015; Zhang *et al*, 2016). Calcineurin can dephosphorylate transcription factors belonging to the MiT-TFE family, TFEB and TFE3, enabling their nuclear translocation and induced transcription of target genes that stimulates biogenesis of lysosomes and autophagosomes (Martina *et al*, 2016; Medina **et al**, 2015; Zhang **et al**, 2016). Another example of adjusting cellular metabolism due to lysosomal dysfunction is activation of transcription factors inducing the expression of genes responsible for cholesterol biosynthesis (Luo *et al*, 2020). It occurs in response to inefficient efflux of cholesterol from lysosomes that leads to its impaired delivery to the endoplasmic reticulum (ER) (Luo **et al**, 2020). Recently, abnormal cholesterol accumulation in lysosomes was shown to increase a cytosolic pool of Ca^2+^ (Tiscione *et al*, 2019) and to promote nuclear accumulation of MiT-TFE factors (Contreras *et al*, 2020; Willett *et al*, 2017).

Lysosomes are the main intracellular compartment where the turnover of membrane proteins occurs (Trivedi *et al*, 2020). The majority of these proteins are delivered to lysosomes by means of endosomal trafficking via a sorting mechanism facilitated by endosomal sorting complexes required for transport (ESCRTs). ESCRTs encompass several protein assemblies (ESCRT-0, I, II, III) that mediate membrane remodeling processes in endocytosis, autophagy, cytokinesis, nuclear envelope sealing and virus budding (Vietri *et al*, 2020). During endosomal sorting, ESCRTs act sequentially to incorporate membrane proteins marked for degradation by ubiquitin into the lumen of endocytic organelles called late or multivesicular endosomes (Raiborg & Stenmark, 2009; Wenzel *et al*, 2018). This occurs by invagination and scission of the endosomal outer (limiting) membrane, thereby forming intraluminal vesicles. Upon fusion of late endosomes with lysosomes, intraluminal vesicles and their cargo reach the lysosomal lumen to be degraded. Inhibition of ESCRT-dependent cargo sorting on endosomes causes accumulation of membrane proteins, such as plasma membrane receptors, on the endosomal limiting membranes, that may activate intracellular signaling pathways (Szymanska *et al*, 2017). As we previously showed, depletion of core ESCRT-I subunits, Vps28 or Tsg101, induces NF-κB signaling initiated by cytokine receptor clustering on endosomal structures (Banach-Orlowska *et al*, 2018; Maminska *et al*, 2016).

Recent reports uncovered a new role of ESCRT complexes, including ESCRT-I, in the turnover of proteins at yeast vacuolar membranes (Morshed *et al*, 2020; Yang *et al*, 2021; Yang *et al*, 2020; Zhu *et al*, 2017). However, it remained unknown whether a similar function could be performed by mammalian ESCRTs at lysosomal membranes. Up to date, mammalian ESCRT proteins have been shown to associate with lysosomes only to restore their integrity disrupted by damaging agents (Jia *et al*, 2020; Radulovic *et al*, 2018; Skowyra *et al*, 2018). Yet, whether ESCRTs are implicated in the physiological regulation of lysosomal morphology, function or signaling in the absence of induced lysosomal damage has not been thoroughly addressed. This question is important for human health, as lysosomal function and signaling contribute to development or progression of cancer (Machado *et al*, 2021; Tang *et al*, 2020), including colorectal cancer (CRC) in which ESCRT machinery has been proposed as a promising therapeutic target (Kolmus *et al*, 2021; Szymanska *et al*, 2020).

Here we show that the ESCRT-I complex sorts membrane proteins at the lysosomes and regulates lysosomal size and function in human colorectal cancer (CRC) cells. Consistently, ESCRT-I deficiency activates transcriptional responses indicative of lysosomal dysfunction including Ca^2+^- and MCOLN1-dependent but cholesterol-independent TFEB/TFE3 signaling. The mechanism of inducing TFEB/TFE3 signaling upon lysosomal stress differs from the one that maintains basal activation of this pathway, which instead of Ca^2+^ and MCOLN1 involves the availability of exogenous lipids.

## Results

### ESCRT-I limits lysosome size

To address whether ESCRT-I is involved in lysosomal membrane homeostasis and signaling we used CRC cells, in which we previously reported ESCRT-I to regulate cell growth and intracellular signaling (Kolmus **et al**, 2021). Using siRNA-mediated approach, we efficiently depleted key ESCRT-I subunits, Tsg101 or Vps28, in RKO cells (**Fig. 1A**). Of note, the removal of one of them reduced the levels of the other due to the ESCRT-I complex destabilization, as described (Bache *et al*, 2004; Kolmus **et al**, 2021). First, we analyzed by confocal microscopy whether knockdown of Tsg101 or Vps28 in RKO cells affected the intracellular distribution of LAMP1, a marker of late endosomes and lysosomes (**Fig. S1**). We noticed that ESCRT-I deficiency increased the area of LAMP1-positive vesicular structures (**Fig. 1B and S1**), suggesting that late endosomes and/or lysosomes were enlarged.

**Fig. 1.**
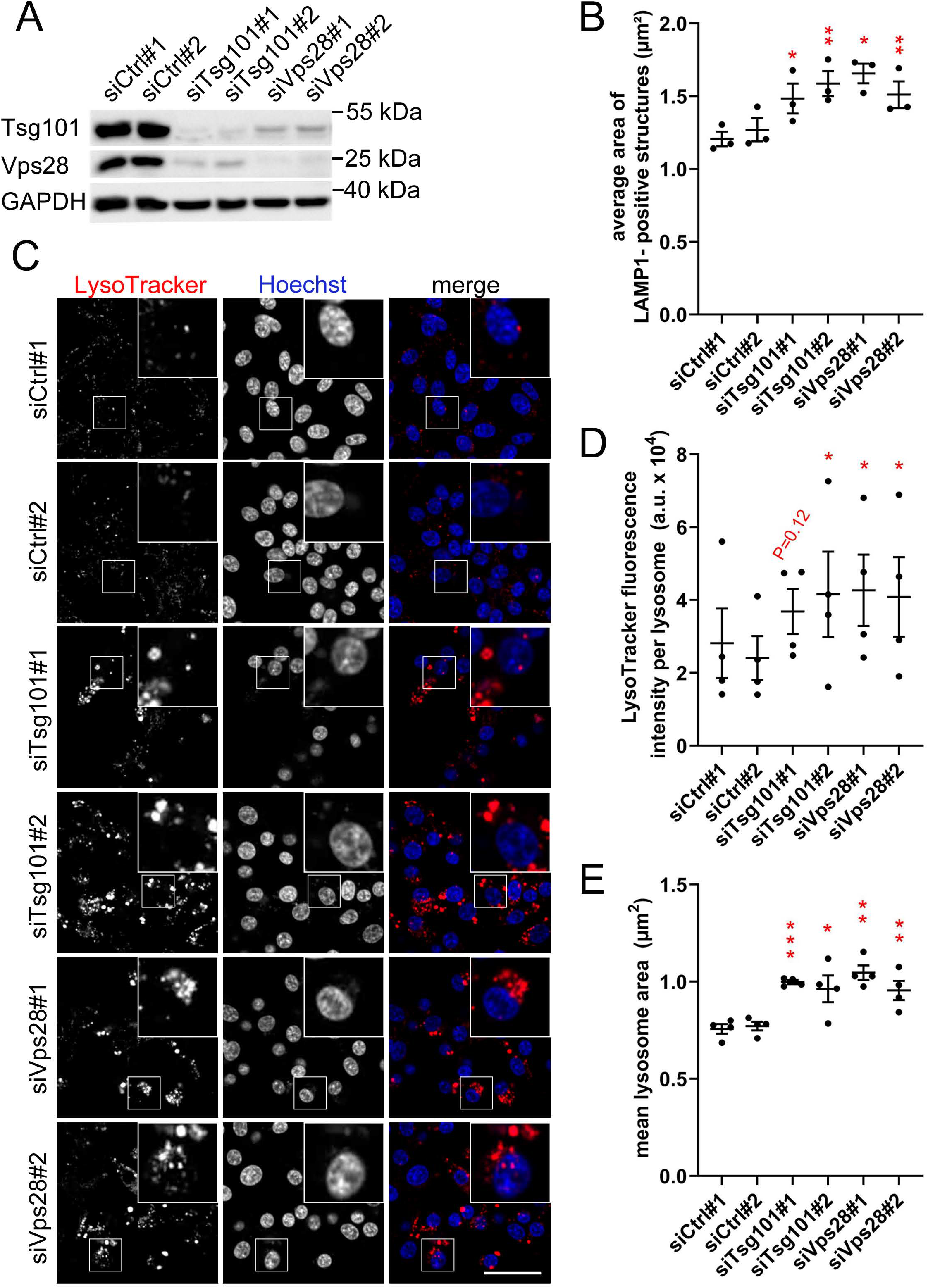
ESCRT-I dysfunction causes enlargement of lysosomes in RKO cells. **(A)** Western blots showing the depletion efficiencies of ESCRT-I subunits, Tsg101 or Vps28 (using two single siRNAs for each subunit, #1 or #2), as compared to control conditions (non-targeting siRNAs, Ctrl#1 or #2), in RKO cells. GAPDH was used as a loading control. **(B)** Dot plot showing the average area of LAMP1-positive vesicles in control or ESCRT-I-depleted cells, calculated based on confocal microscopy images (shown in **Supplementary Fig. 1**). Values derived from independent experiments (dots) and their means (n=3 +/− SEM) are presented. Statistical significance tested by comparison to averaged values measured for siCtrl#1 and #2 *P < 0.05, **P<0.01. **(C)** Maximum intensity projection confocal images of live cells, showing the intracellular distribution of lysosomes stained with LysoTracker dye (red) in control cells and upon ESCRT-I depletion. Cell nuclei marked with Hoechst stain (blue). Scale bar, 50 μm. **(D-E)**. Dot plots showing average fluorescence intensity of LysoTracker expressed in arbitrary units (a.u., **D**) and average area (**E**) of detected lysosomal structures in control or ESCRT-depleted cells, calculated based on live cell microscopy images (shown in C). Values derived from independent experiments (dots) and their means (n=4 +/− SEM) are presented. Statistical significance tested by comparison to averaged values measured for siCtrl#1 and #2. *P<0.05, **P<0.01, ***P<0.001.

To confirm the enlargement of lysosomes in the absence of ESCRT-I, we measured the effect of Tsg101 or Vps28 depletion on the intracellular distribution of LysoTracker, a cell-permeable fluorescent dye that accumulates in non-damaged acidic lysosomes (Pierzynska-Mach *et al*, 2014) (**Fig. 1C**). Lack of ESCRT-I augmented the LysoTracker staining intensity per lysosome and increased the size of lysosomes (**Fig. 1D–E**). Strong LysoTracker accumulation in lysosomes upon ESCRT-I deficiency indicated that their integrity was not impaired and that they maintained an acidic pH, important for their degradative function (Ballabio, 2016). To verify that the observed effects of ESCRT-I depletion on lysosomal size are not specific only to RKO cells, we depleted Tsg101 or Vps28 in another CRC cell line, DLD-1 (**Fig. S2A**). Reassuringly, we again observed increased LysoTracker staining intensity and size of lysosomes (**Fig. S2B-D**).

Cumulatively, we discovered that in CRC cells, ESCRT-I limits the size of lysosomes.

### ESCRT-I mediates the turnover of lysosomal membrane proteins, including MCOLN1

We investigated the mechanism through which ESCRT-I controls lysosomal size. As recently shown, degradation of proteins residing in the lysosomal membrane requires their internalization into the lumen of lysosomes together with adjacent membrane parts (Lee *et al*, 2020). Hence, we hypothesized that the enlargement of lysosomes in the absence of ESCRT-I could be caused by inhibited internalization of parts of lysosomal membranes due to impaired turnover of resident membrane proteins. To address this, we analyzed by confocal microscopy the amount of ubiquitinated proteins on lysosomes using antibodies recognizing mono- and polyubiquitinated protein conjugates. In control RKO cells, we observed a weak ubiquitin staining on the LysoTracker- and LAMP1-positive structures, likely reflecting ubiquitinated cargo targeted for lysosomal degradation (**Fig. 2A**). However, depletion of ESCRT-I subunits led to a strong accumulation of ubiquitin on vesicular structures. This included enlarged lysosomes, marked by LysoTracker staining, in which ubiquitin was particularly enriched on their membranes marked by LAMP1 (**Fig. 2A**). This pointed to an impaired degradation of lysosomal membrane proteins.

**Fig. 2.**
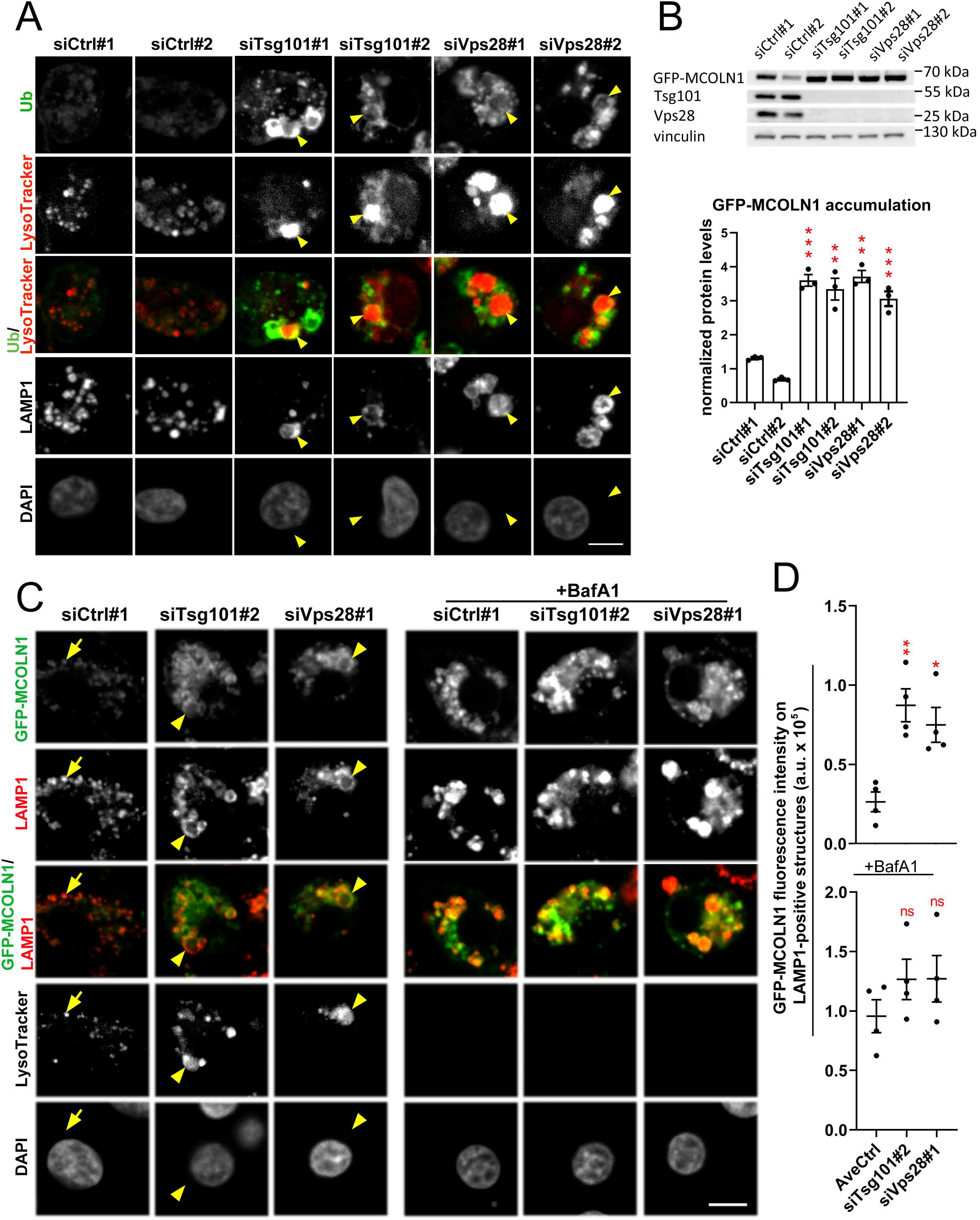
ESCRT-I mediates the degradation of lysosomal membrane proteins in RKO cells. **(A)** Representative maximum intensity projection confocal images of fixed RKO cells showing intracellular distribution of ubiquitin (green), LysoTracker dye (red) and LAMP1 (grey) in ESCRT-I-depleted (siTsg101#1 or #2, siVps28#1 or #2) or control (Ctrl#1, #2, non-targeting siRNAs) cells. Cell nuclei marked with DAPI stain (grey). Enlarged lysosomes enriched in ubiquitinated proteins at lysosomal outer membranes, marked with LAMP1, are indicated by arrowheads. Scale bar, 10 μm. **(B)** Representative western blot (upper panel) showing the levels of ectopically expressed GFP-MCOLN1 and ESCRT-I components in control or ESCRT-I-depleted RKO cells. The graph (lower panel) shows GFP-MCOLN1 levels expressed as fold change with respect to averaged values measured for siCtrl#1 and #2 by densitometry analysis of western blotting bands. Vinculin was used as a gel loading control. Values derived from independent experiments and their means (n=4 +/− SEM) are presented. Statistical significance tested by comparison to siCtrl#1. **P<0.01, ***P<0.001. **(C)** Representative single confocal plane images of fixed cells showing the effect of ESCRT-I depletion and/or 18 h bafilomycin A1 (BafA1) treatment on the intracellular distribution of ectopically expressed GFP-MCOLN1 (green), with respect to LAMP1 (red) and LysoTracker dye (grey). In control cells, GFP-MCOLN1 accumulation on LAMP1-positive vesicles indicated by arrows. GFP-MCOLN1 accumulation on enlarged LAMP1-positive lysosomal structures in ESCRT-I-depleted cells indicated by arrowheads. Cell nuclei marked with DAPI stain (grey). Scale bar, 10 μm. **(D)** Dot plot showing the fluorescence intensity of GFP-MCOLN1 colocalizing with LAMP1-positive vesicles expressed in arbitrary units (a.u.) in confocal microscopy images of control or ESCRT-I-depleted cells and/or upon18 h bafilomycin A1 (BafA1) treatment (shown in C). Values derived from independent experiments (dots) and their means (n=4 +/− SEM) are presented. Statistical significance tested by comparison to averaged values measured for siCtrl#1 and #2 (AveCtrl). ns—non-significant (P≥0.05), **P<0.01.

To verify the inhibited turnover of proteins from lysosomal membranes upon ESCRT-I depletion, we generated an RKO cell line stably expressing GFP-tagged mucolipin 1 (GFP-MCOLN1, **Fig. 2B**), a Ca^2+^ channel protein recently shown to be removed from lysosomal membranes by its internalization into the lumen (Lee **et al**, 2020). As expected (Cheng *et al*, 2010), we observed by confocal microscopy that in control cells, GFP-MCOLN1 localized predominantly to LAMP1-positive structures, both LysoTracker-negative late endosomes and LysoTracker-positive lysosomes **(Fig. 2C)**. Knockdown of ESCRT-I subunits increased the levels of GFP-MCOLN1 protein, observed by western blotting and microscopy (**Fig. 2B–C**). Importantly, upon ESCRT-I depletion, GFP-MCOLN1 strongly accumulated on LAMP1-positive structures (**Fig. 2C–D**), including enlarged lysosomes (**Fig. 2C**). To address whether the observed accumulation of GFP-MCOLN1 occurred due to impaired lysosomal degradation, we used bafilomycin A1 (BafA1), an inhibitor of the V-ATPase proton pump (Yoshimori *et al*, 1991). Consistent with its inhibitory effect on lysosomal acidification, BafA1 led to a loss of LysoTracker staining in both control and ESCRT-I-depleted cells (**Fig. 2C**). In control cells, BafA1 caused a strong accumulation of GFP-MCOLN1 protein on LAMP1-positive structures (**Fig. 2C–D**). However, it did not significantly increase the amount of GFP-MCOLN1 already enriched on LAMP1-positive structures in cells lacking Tsg101 or Vps28 (**Fig. 2C–D**). These results indicate that GFP-MCOLN1 is constantly degraded in lysosomes, however ESCRT-I depletion inhibits its degradation and thereby causes its accumulation on lysosomal membranes.

Altogether, our data argue that ESCRT-I mediates the lysosomal degradation of late endosomal and lysosomal membrane proteins, including MCOLN1. This finding corroborated our hypothesis that the enlargement of lysosomes in CRC cells lacking ESCRT-I could stem from inhibited turnover of lysosomal membrane proteins.

### ESCRT-I deficiency activates transcriptional responses indicative of lysosomal stress

We reasoned that the inhibited turnover of membrane proteins from the surfaces of lysosomes in the absence of ESCRT-I could impair the function of these organelles. Lysosomal dysfunction is often reflected by the activation of specific transcriptional responses (Di Malta *et al*, 2019; Zhao *et al*, 2020). Thus, we performed RNA-sequencing to investigate whether ESCRT-I depletion in RKO cells leads to changes in gene expression that would point to altered lysosomal function. Gene ontology (GO) analysis of commonly upregulated genes after Tsg101 or Vps28 depletion identified an elevated inflammatory response, consistent with our previous reports (Banach-Orlowska **et al**, 2018; Maminska **et al**, 2016). Importantly, we also observed an enhanced expression of genes annotated to autophagy and cholesterol metabolism (**Fig. S3A**), two processes highly dependent on lysosomes (Ikonen, 2008; Lieberman *et al*, 2012; Medina **et al**, 2015; Sardiello *et al*, 2009).

Among genes annotated to autophagy, whose expression was induced upon ESCRT-I depletion, we identified a group of established MiT-TFE targets involved in lysosomal biogenesis and autolysosomal degradation (**Fig. 3A**) (Medina **et al**, 2015; Perera & Zoncu, 2016; Settembre *et al*, 2012). Thus, we tested whether the localization of TFEB and TFE3 transcription factors was affected by ESCRT-I deficiency. By western blotting analysis, we observed that depletion of Tsg101 or Vps28 increased the abundance of both TFEB and TFE3 in whole-cell lysates as well as nuclear fractions of RKO cells (**Fig. 3B**). Next, we verified whether the induced expression of genes involved in the regulation of lysosomal function in ESCRT-I-depleted RKO cells occurred due to the MiT-TFE activation. To this end, we depleted TFEB and TFE3 factors using siRNA. We observed that their simultaneous depletion in cells lacking Tsg101 or Vps28 prevented the elevated expression of two MiT-TFE target genes that encode lysosomal proteins, NPC1 and MCOLN1 (**Fig. 3C**).

**Fig. 3.**
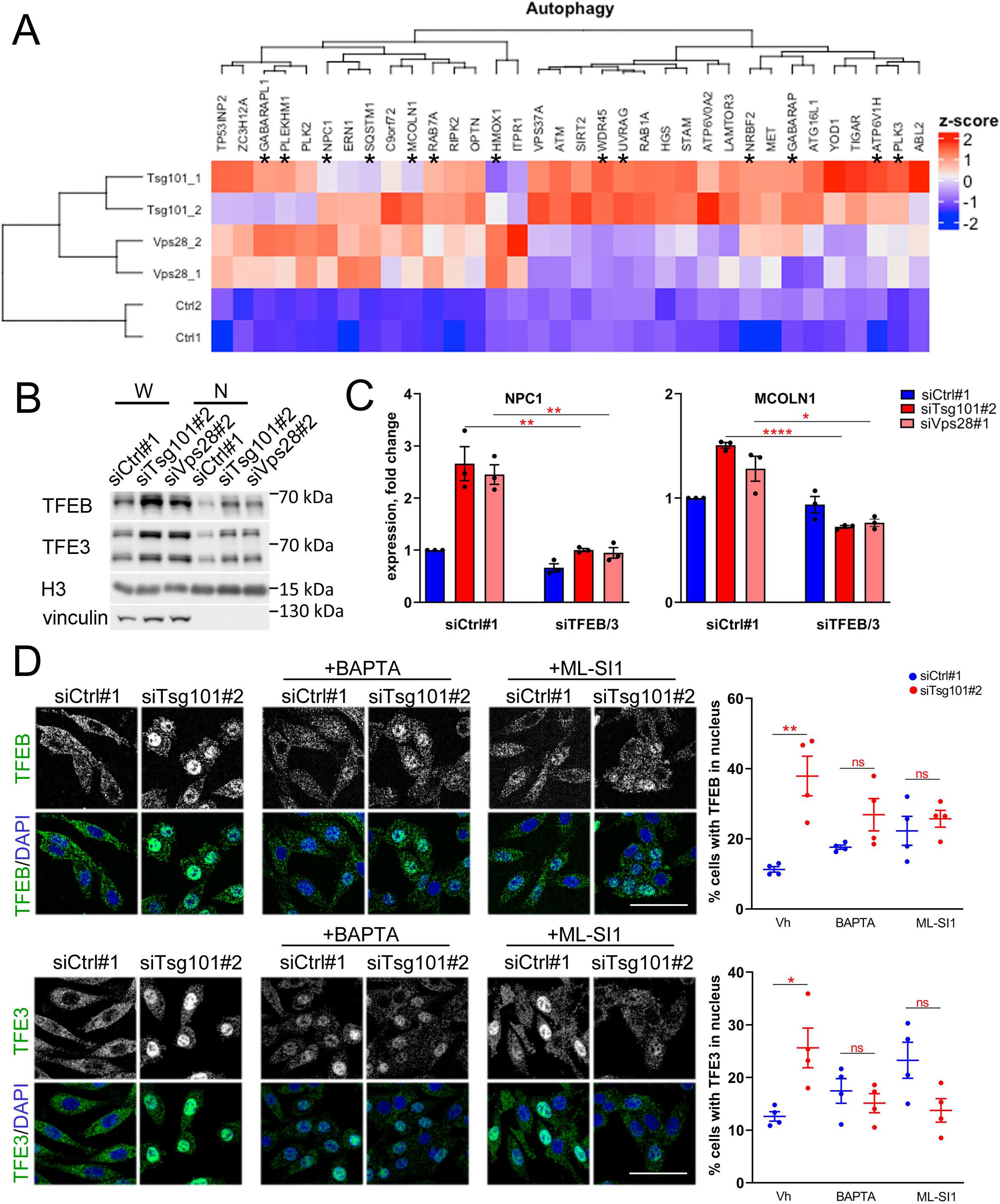
Depletion of ESCRT-I induces Ca^2+^/MCOLN1-dependent TFEB/TFE3 signaling. **(A)** Heatmap visualizing expression of genes annotated to “autophagy” gene ontology (GO:0006914) process, whose mRNA levels were detected by RNA-seq as upregulated after Tsg101 or Vps28 depletion in RKO cells. Established MiT-TFE target genes are indicated by asterisks. **(B)** Western blots showing levels of TFEB and TFE3 proteins in whole-cell lysates (W) and nuclear fractions (N) of RKO cells. To examine the fraction purity, the levels of vinculin (cytosolic marker) were probed. H3 protein was used as a loading control for nuclear fractions. **(C)** qPCR results showing the expression of MiT-TFE target genes upon ESCRT-I and/or MiT-TFE depletion (using single siRNAs for Tsg101, Vps28, TFEB or TFE3) presented as fold changes with respect to control cells (treated with non-targeting siRNA, Ctrl#1). Values derived from independent experiments and their means (n=4 +/− SEM) are presented. Statistical significance tested by comparison to siTsg101 or siVps28 conditions. *P < 0.05, **P<0.01, ****P<0.0001. **(D)** Representative maximum intensity projection confocal images (left panels) of fixed RKO cells 48 hpt showing intracellular distribution of TFEB or TFE3 (green) in ESCRT-I-depleted (siTsg101#2) or control (Ctrl#1, non-targeting siRNA) cells and/or upon 2 h treatment with BAPTA-AM (10 μM) or ML-SI1 (25 μM). Cell nuclei marked with DAPI stain (blue). Scale bar, 50 μm. Dot plots (right panels) showing the percentage of cells with nuclear TFEB or TFE3 localization. Values derived from independent experiments (dots) and their means (n=4 +/− SEM) are presented. Statistical significance tested by comparison to siCtrl#1 conditions. ns—non-significant (P≥0.05), *P <0.05, **P<0.01.

Hence, ESCRT-I depletion leads to a transcriptional response to lysosomal stress. This includes elevated expression of genes encoding lysosomal proteins, due to the activation of TFEB and TFE3 transcription factors.

### Depletion of ESCRT-I activates TFEB and TFE3 transcription factors in an MCOLN1-dependent manner

Although our results revealing elevated MiT-TFE signaling upon ESCRT-I depletion (**Fig. 3A–C**) were obtained 72 h post transfection (hpt) of cells with siRNA, we suspected that activation of these transcription factors could occur earlier. To this end, by quantitative analysis of confocal microscopy images, we measured the percentage of RKO cells with TFEB or TFE3 present in their nuclei. We detected nuclear TFEB or TFE3 in around 10% of control cells, a fraction that was constant at different time-points post transfection (**Fig. S3B**). However, upon Tsg101 depletion, the percentage of RKO cells with nuclear accumulation of these transcription factors increased significantly already at 48 hpt, in the case of TFEB to the similar level (around 30%) as observed at 72 hpt (**Fig. S3B**).

As shown above, in cells lacking ESCRT-I, the impaired turnover of MCOLN1, a Ca^2+^ channel, was associated with the activation of MiT-TFE signaling. Nuclear translocation of TFEB or TFE3 may occur upon their dephosphorylation by calcineurin, a calcium-dependent phosphatase (Martina **et al**, 2016; Medina **et al**, 2015; Zhang **et al**, 2016). Moreover, TFEB may be activated by Ca^2+^ exported from lysosomes via MCOLN1 channel (Medina **et al**, 2015; Zhang **et al**, 2016). Hence, we tested whether MCOLN1-dependent Ca^2+^ transport could activate TFEB or TFE3 upon ESCRT-I depletion, by applying BAPTA-AM, a chelator of intracellular Ca^2+^ (Medina **et al**, 2015), or ML-SI1, an inhibitor of MCOLN1 channel activity (Sun *et al*, 2018). To elucidate mechanisms that induce the MiT-TFE signaling upon ESCRT-I depletion, we applied these compounds at an early stage of the pathway activation (48 hpt). Surprisingly, BAPTA-AM or ML-SI1 treatment did not decrease the nuclear levels of TFEB and TFE3 in control cells, indicating that basal activation of MiT-TFE factors is not mediated by Ca^2+^ signaling (**Fig. 3D**). Nevertheless, Ca^2+^ chelation or MCOLN1 inhibition prevented the nuclear accumulation of TFEB and TFE3 proteins in cells lacking Tsg101 (**Fig. 3D**).

Taken together, these results showed that ESCRT-I deficiency leads to the prolonged activation of MiT-TFE signaling that is initiated in a Ca^2+^- and MCOLN1-dependent manner.

### Depletion of ESCRT-I inhibits lysosomal cholesterol efflux that does not contribute to activation of TFEB/TFE3 signaling

Our data confirming the activation of TFEB and TFE3 signaling in ESCRT-I-depleted cells reinforced our hypothesis that ESCRT-I deficiency could lead to lysosomal dysfunction. TFEB activation may occur upon inhibition of cholesterol efflux from lysosomes (Boutry *et al*, 2019; Willett **et al**, 2017). Consistent with this, ESCRT-I depletion led to elevated expression of genes encoding enzymes of cholesterol biosynthesis (**Fig. 4A**), suggestive of impaired delivery of cholesterol from lysosomes to the ER (Luo **et al**, 2020; Xue *et al*, 2020). In accordance with this, we observed that ESCRT-I depletion in RKO cells led to the accumulation of cholesterol (stained by filipin) in enlarged LAMP1-positive structures (**Fig. 4B, S4A**). This accumulation was suppressed when cells were cultured in a delipidated medium (**Fig. 4B, S4A**) that has reduced levels of lipids, including cholesterol which under such conditions is not delivered to lysosomes via endocytic uptake (Brovkovych *et al*, 2019). These data argue that cholesterol accumulated in lysosomes upon ESCRT-I depletion is of extracellular origin.

**Fig. 4.**
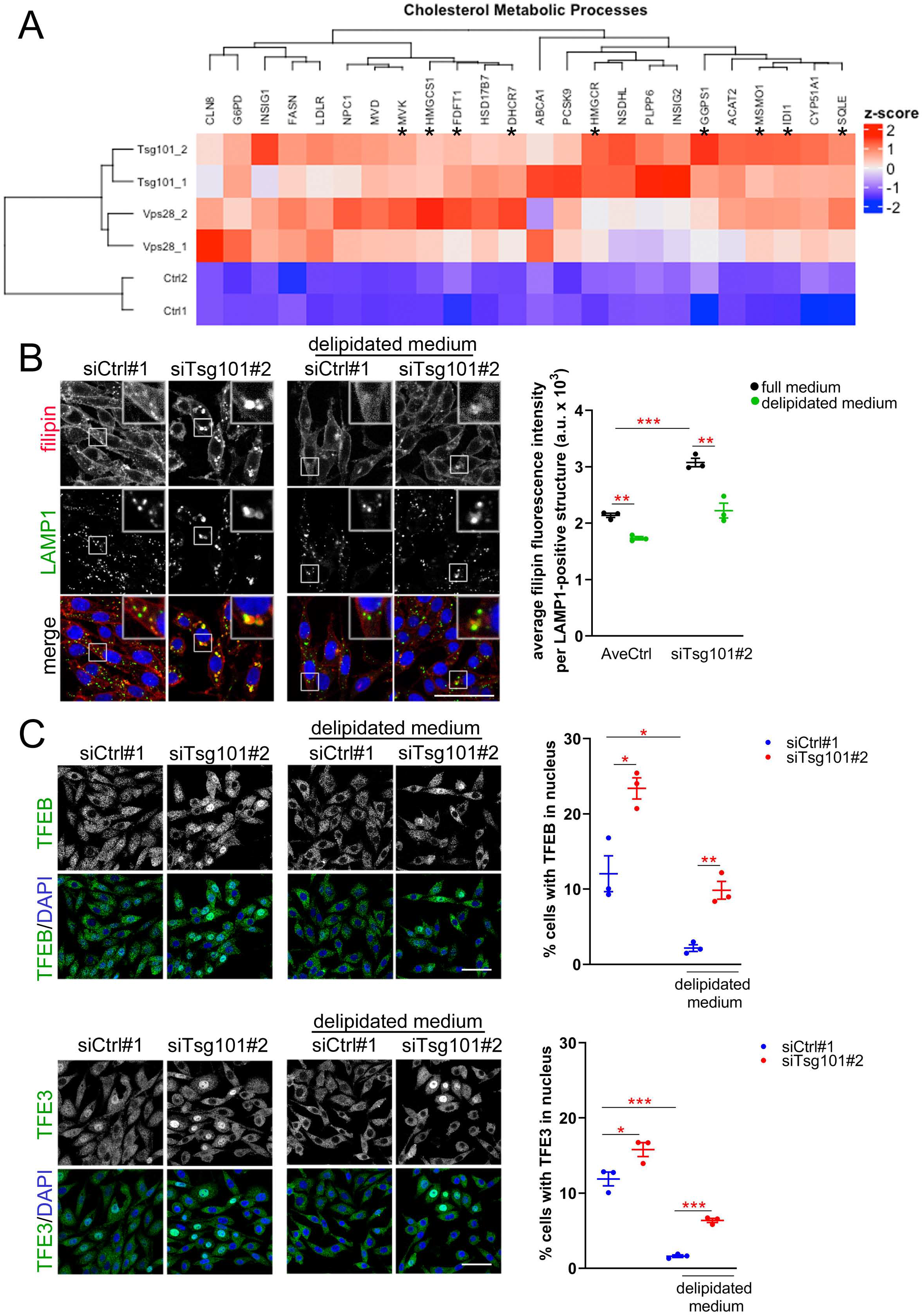
ESCRT-I deficiency inhibits cholesterol efflux from lysosomes that does not contribute to nuclear accumulation of TFEB and TFE3 factors. **(A)** Heatmap visualizing expression of genes annotated to “cholesterol metabolic processes” gene ontology (GO:0008203) process, whose mRNA levels were detected by RNA-seq as upregulated after Tsg101 or Vps28 depletion in RKO cells. Established cholesterol biosynthesis genes are indicated by asterisks. **(B)** Representative maximum intensity projection confocal images of fixed RKO cells 48 hpt (left panel) showing the intracellular distribution of free cholesterol marked with filipin dye (red) with respect to LAMP1 (green) expressed in arbitrary units (a.u.) in control or Tsg101-depleted cells and/or upon 40 h culture in delipidated medium. Cell nuclei marked with DRAQ7 dye (blue). Scale bar, 50 μm. Dot plot (right panel) showing the average filipin fluorescence intensity per LAMP1-positive structure. Values derived from independent experiments (dots) and their means (n=3 +/− SEM) are presented. Statistical significance tested by comparison to averaged values measured for control cells (AveCtrl) and/or siTsg101#2 conditions. **P<0.01, ***P<0.001. **(C)** Representative maximum intensity projection confocal images of fixed RKO cells 48 hpt (left panels) showing intracellular distribution of TFEB or TFE3 (green) in ESCRT-I-depleted (siTsg101#2) or control (Ctrl#1, non-targeting siRNA) cells and/or upon 40 h culture in delipidated medium. Dot plots (right panels) showing the percentage of cells with nuclear TFEB or TFE3 localization. Values derived from independent experiments (dots) and their means (n=3 +/− SEM) are presented. Statistical significance tested by comparison to siCtrl#1 conditions. *P <0.05, **P<0.01, ***P<0.001.

To verify whether the upregulated expression of cholesterol biosynthesis genes in the absence of ESCRT-I is due to an impaired efflux of lysosomal cholesterol to the ER, we supplied cells with cholesterol in a soluble form that reaches the ER independently of the endolysosomal trafficking (Trinh *et al*, 2020). As analyzed by quantitative RT-PCR, soluble cholesterol supplementation prevented the upregulation of cholesterol biosynthesis genes in RKO cells lacking Tsg101 or Vps28 (**Fig. S4B**). This confirmed an impaired delivery of cholesterol from lysosomes to the ER in the absence of ESCRT-I.

Impaired cholesterol efflux from lysosomes can cause the import of extracellular calcium to the cytoplasm (Boutry **et al**, 2019; Tiscione **et al**, 2019) that may lead to TFEB nuclear translocation (Boutry **et al**, 2019). Hence, we tested whether preventing the lysosomal cholesterol accumulation by culturing cells in a delipidated medium (as shown in **Fig. 4B**) affected the induction of TFEB/TFE3 signaling. Intriguingly, we observed that deprivation of exogenous lipids strongly reduced the percentage of control cells with nuclear TFEB or TFE3 (**Fig. 4C**), indicating that basal activation of this pathway depends on lipid delivery. However, culture of cells in the absence of exogenous lipids did not prevent the increase in the number of cells with nuclear TFEB or TFE3 upon Tsg101 depletion.

Collectively, we show that in the absence of ESCRT-I, lysosomes are not fully functional as they are defective in the export of cholesterol, which is reflected by induced transcription of cholesterol biosynthesis genes. However, although the availability of exogenous lipids contributes to the activation of TFEB/TFE3 signaling under basal growth conditions, the accumulation of cholesterol in lysosomes upon ESCRT-I deficiency is not a causative factor for the induction of this pathway. Based on these findings and on differential effects of inhibiting the Ca^2+^-MCOLN1 pathway on the nuclear abundance of TFEB and TFE3 (shown in **Fig. 3D**), we conclude that distinct mechanisms underlie TFEB/TFE3 signaling during normal growth or upon lysosomal stress.

## Discussion

Lysosomes have recently emerged as important players in cancer development or progression, while targeting their function has been proposed as a promising strategy in cancer treatment (Machado **et al**, 2021; Tang **et al**, 2020). Recently, we showed that components of the ESCRT machinery could serve as potential therapeutic targets against colorectal cancer (CRC) (Kolmus **et al**, 2021; Szymanska **et al**, 2020), but whether modulation of ESCRT activity would affect lysosomal function in CRC cells has not been tested. Here, by a comprehensive analysis of the morphology and function of lysosomes in CRC cells, we discover that mammalian ESCRT-I proteins, Tsg101 or Vps28, control lysosomal size and homeostasis, regulating lysosomal membrane protein turnover and cholesterol efflux. Due to the involvement of ESCRT-I in lysosomal homeostasis, the absence of this complex activates transcriptional responses related to biogenesis of lysosomes and cholesterol biosynthesis.

Although ESCRT-dependent degradation of vacuolar transmembrane proteins has been shown to occur in yeast (Morshed **et al**, 2020; Yang **et al**, 2021; Yang **et al**, 2020; Zhu **et al**, 2017), an analogous mechanism has not been fully characterized in mammalian cells. So far, mammalian ESCRT proteins were shown to associate with lysosomes to repair their membranes only upon damage (Eriksson *et al*, 2020; Radulovic **et al**, 2018; Skowyra **et al**, 2018). Our study extends these findings by showing that ESCRT-I has a broader function as it maintains lysosomal homeostasis. This is consistent with a very recent discovery of Zhang **et al** who uncovered that ESCRT components, particularly ESCRT-III and Vps4, are involved in degradation of several lysosomal membrane proteins in HEK293 and HeLa cells (Zhang **et al**, 2021).

Previous analyses of LAMP1-positive compartment suggested that depletion of ESCRT subunits might affect lysosomal morphology (Doyotte *et al*, 2005; Du *et al*, 2012; Du *et al*, 2013; Szymanska **et al**, 2020). However, as LAMP1 is a marker of both late endosomes and lysosomes (Saftig & Klumperman, 2009), these observations did not allow conclusions about the status of lysosomes. Here, by investigating the LysoTracker-positive compartment, we establish that ESCRT-I depletion leads to the enlargement of lysosomes due to impaired degradation of lysosomal membrane proteins, including MCOLN1. Our study not only confirms that MCOLN1 is degraded from the lysosomal surface as recently reported (Lee **et al**, 2020) but also unravels that this degradation is mediated by ESCRT-I.

MCOLN1 protein has been shown to promote nuclear translocation of transcription factor TFEB, through activation of calcineurin phosphatase by Ca^2+^ released from lysosomes (Medina **et al**, 2015). Calcineurin-dependent activation of TFE3 was also reported (Martina **et al**, 2016) but whether it is mediated by MCOLN1 has not been studied. We show that upon ESCRT-I depletion, both TFEB and TFE3 are activated by Ca^2+^ in an MCOLN1-dependent manner, clarifying that the two factors share a similar mechanism of activation upon lysosomal dysfunction. Importantly, in cells lacking ESCRT-I, the nuclear translocation of these transcription factors is associated with an elevated abundance of non-degraded MCOLN1 on enlarged lysosomes. Therefore we propose that accumulation of MCOLN1 may serve as a mechanism of TFEB/TFE3 induction from lysosomes with an inhibited turnover of their membrane proteins.

The initial discoveries concerning activation of TFEB focused on conditions, such as starvation or inhibitor treatments, applied for relatively short time, up to few hours (Medina **et al**, 2015; Settembre *et al*, 2011; Settembre **et al**, 2012; Zhang **et al**, 2016). We reveal that ESCRT-I depletion leads to a prolonged activation of TFEB and TFE3 signaling, as it has been shown to occur due to cellular stress, such as pathogen infection (Pastore *et al*, 2016) or ER stress (Martina **et al**, 2016). Previously, we discovered that dysfunction of endolysosomal trafficking due to ESCRT-I deficiency activates NF-κB-mediated stress response, driven by non-degraded cytokine receptors accumulated on endosomes (Banach-Orlowska **et al**, 2018; Maminska **et al**, 2016). Here, we discover another stress response pathway activated upon ESCRT-I depletion, namely TFEB/TFE3 signaling that is likely induced as a consequence of disrupted lysosomal homeostasis. Whether ESCRT-I restricts the TFEB/TFE3 pathway by facilitating the turnover of lysosomal membrane proteins remains to be confirmed.

As reported, depletion of some components of ESCRT complexes, namely Hrs (ESCRT-0 subunit) or Vps4A/B cause accumulation of cholesterol on LAMP1-positive structures (Du **et al**, 2012; Du **et al**, 2013). These studies, based on experiments performed in HeLa cells, concluded that Tsg101 or ESCRT-III subunits are not involved in cholesterol transport from lysosomes, arguing that the roles of Hrs or Vps4A/B in this process are independent of other ESCRTs. However, our finding that the disruption of ESCRT-I complex by depletion of Tsg101 or Vps28 leads to accumulation of cholesterol in lysosomes of RKO cells argues that lysosomal cholesterol efflux requires functional ESCRT machinery. The discrepancy between our results and published data (Du **et al**, 2012) regarding the effect of Tsg101 depletion may be due to cell type-specific effects.

Aberrant cholesterol transport from lysosomes has been associated with increased cytoplasmic Ca^2+^ levels (Tiscione **et al**, 2019) and MiT-TFE activation (Boutry **et al**, 2019; Contreras **et al**, 2020; Willett **et al**, 2017). Although we observed a similar association in cells lacking ESCRT-I, we could not confirm a causal relationship between improper intracellular cholesterol distribution and activation of TFEB/TFE3 signaling. Cancer cells often have constitutive activation of transcription factors from the MiT-TFE family which improves lysosomal degradation to support accelerated rates of growth (Perera *et al*, 2019). Intriguingly, our data show that basal TFEB/TFE3 nuclear accumulation in RKO cells is not mediated by Ca^2+^ but depends on the presence of lipids in the medium. Therefore we identify two different ways to activate TFEB/TFE3 signaling in cancer cells: lipid-dependent basal activation or stress-induced, MCOLN1- and Ca^2+^-dependent activation.

Hence, our work provides insights into the interplay between basal and stress-adaptive roles of TFEB and TFE3 transcription factors. Understanding such interplay may point to new strategies for cancer treatment (Perera **et al**, 2019). Further studies may address whether cancer cells with increased basal lysosomal function via MiT-TFE signaling could be particularly dependent on lysosomal membrane protein turnover that, as we show here, is mediated by ESCRT-I.

## Materials and Methods

### Antibodies

The following antibodies were used: anti-Tsg101 (Cat# ab83), anti-Vps28 (Cat# ab167172) from Abcam; anti- TFEB (Cat # 4240S), anti-TFE3 (Cat# 14779S) from Cell Signaling Technologies; anti-LAMP1 (Cat# H4A3) from DSHB; anti-mono- and -polyubiquitinylated conjugates (Cat# BML-PW8810) from Enzo Life Sciences; anti-GFP (Cat# AF4240) from R&D Systems; anti-GAPDH (Cat# sc-25778) from Santa Cruz; anti-histone H3 (Cat# H0164), anti-LAMP1 (Cat# L-1418), anti-vinculin (Cat# V9131-.2ML) from Sigma; secondary horseradish peroxidase (HRP)-conjugated goat anti-mouse, goat anti-rabbit, and bovine anti-goat antibodies (Jackson ImmunoResearch); secondary Alexa Fluor 488-conjugated donkey anti-mouse and Alexa Fluor 647-conjugated donkey anti-rabbit antibodies from Thermo Fisher Scientific.

### Plasmids

To obtain EGFP-MCOLN1 construct for lentiviral transduction, human MCOLN1 was amplified by PCR with the following oligonucleotides with overhangs (underlined): 5’-<u>GACACCGACTCTAGA</u>ATGGTGAGCAAGGGCGAGGAGC-3’ (forward) and 5’-<u>AACTAGTCCGGATCC</u>TCAATTCACCAGCAGCGAATGC-3’ (reverse) from Mucolipin1-pEGFP C3 (Addgene plasmid #62960) construct and subcloned into XbaI and BamHI restriction sites of pLenti-CMV-MCS-GFP-SV-puro (Addgene plasmid #73582) vector using sequence- and ligation-independent cloning (SLIC) method as described elsewhere (Jeong *et al*, 2012). Mucolipin1-pEGFP C3 (Addgene plasmid#62960) construct was a gift from Paul Luzio (Pryor *et al*, 2006). psPAX2 (Addgene plasmid # 12260) and pMD2.G (Addgene plasmid # 12259) lentiviral packaging plasmids were a gift from Didier Trono.

### Cell culture and treatment

DLD-1 colon adenocarcinoma and HEK293T embryonic kidney cells were maintained in Dulbecco’s modified Eagle’s medium (DMEM, Sigma-Aldrich, M2279) supplemented with 10% (v/v) fetal bovine serum (FBS, Sigma-Aldrich, F7524) and 2 mM L-Glutamine (Sigma-Aldrich, G7513). RKO colon carcinoma cells were cultured in Eagle’s minimum essential medium (EMEM, ATCC, 30-2003) supplemented with 10% (v/v) FBS. Both cell lines were regularly tested as mycoplasma-negative and their identities were confirmed by short tandem repeat (STR) profiling performed by the ATCC Cell Authentication Service.

BAPTA-AM was applied to chelate the intracellular pool of calcium at 10 μM concentration for 2 h. To inhibit MCOLN1 calcium channel activity, ML-SI1 was used for 2 h at 25 μM. Bafilomycin A1 at 50 nM concentration was applied for 18 h to inhibit lysosomal degradation. 20 μM water-soluble cholesterol was added for 48 h. Cells were live-stained with 50 nM or 500 nM LysoTracker for 30 min for live or fixed cell imaging, respectively. EMEM medium supplemented with delipidated FBS (S181 L, Biowest) was used for 40 h to deprive cells of exogenous lipids.

### Cell transfection and lentiviral transduction

RKO cells were seeded on 6-well plates (0.8×10^5^ cells/well) for western blotting and quantitative real-time PCR (qRT-PCR) experiments, on P100 dish (0.9×10^6^ cells per dish) for cellular fractionation experiment or on 0.2% gelatin (G1890, Sigma Aldrich)-covered 96-well plate (Grainer Bio-One, 655-090) (0.25 or 0.4×10^3^ cells/well) for confocal microscopy. DLD-1 cells were seeded on a 6-well plate (0.8×10^5^ cells/well) for western blotting or on 96-well plate (Grainer Bio-One, 655-090) (0.2×10^3^ cells/well) for microscopy. 24 h after seeding, cells were transfected with 30 nM siRNAs using Lipofectamine™ RNAiMAX Transfection Reagent (Thermo Fisher Scientific, 13778150) according the manufacturer’s instructions and imaged or harvested after 48 or 72 h post transfection (hpt). The following Ambion Silencer Select siRNAs (Thermo Fisher Scientific) were used: Negative Control No. 1 (siCtrl#1, 4390843) and Negative Control No. 2 (siCtrl#2, 4390846); siTsg101#1 (s14439), siTsg101#2 (s14440), siVps28#1 (s27577), siVps28#2 (s27579), siTFEB (s15496), siTFE3 (s14031). In experiments with simultaneous knockdown of three genes, the total concentration of siRNA was adjusted to 60 nM using siCtrl#1.

For overexpression of EGFP-tagged MCOLN1, lentiviral particles were produced in HEK293T cells using packaging plasmids: psPAX2 and pMD2.G, as described elsewhere (Barde *et al*, 2010). Subsequently, 1×10^6^ RKO cells were grown in 5 ml of virus-containing EMEM medium on P60 dish for 24 h. Then, cells were split and grown in selection EMEM medium containing 1 μg/ml puromycin for 72 h.

### Lysosomal staining

For live cell imaging, cells were incubated for 30 min with 50 nM LysoTracker as described elsewhere (Hirst *et al*, 2015). Subsequently, cells were washed with probe-free medium and immediately imaged. For fixed cell imaging, cells were incubated for 30 min with 500 nM LysoTracker and fixed with 3.6% paraformaldehyde for 15 min on ice followed by 15 min incubation at room temperature. After three washes with PBS (phosphate buffer saline) cells were immunostained.

### Immunofluorescence staining and microscopy

Cells seeded on 0.2% gelatin-coated plates (Greiner Bio-One, 655-090) were fixed with 3.6% paraformaldehyde at room temperature and immunostained as described elsewhere (Maminska **et al**, 2016). Cell nuclei were marked with DAPI (Sigma-Aldrich, D9542), Hoechst (Thermo Fisher Scientific, Cat no H1399) or DRAQ7 (Thermo Fisher Scientific, Cat no D15106) dye, as indicated in the figure legends. Plates were scanned using Opera Phenix high content screening microscope (PerkinElmer) with 40 × 1.1 NA water immersion objective. Harmony 4.9 software (PerkinElmer) was applied for image acquisition and their quantitative analysis. For quantification of chosen parameters (average area of vesicular structures, fluorescence intensity per structure, percentage of cells with nuclear staining), more than 15 microscopic fields were analyzed per each experimental condition. Maximum intensity projection images were obtained from 3 to 5 z-stack planes with 1 μm interval. Pictures were assembled in ImageJ and Photoshop (Adobe) with only linear adjustments of contrast and brightness.

### Western blotting

Cells were lysed in RIPA buffer (1% Triton X-100, 0.5% sodium deoxycholate, 0.1% SDS, 50 mM Tris pH 7.4, 150 mM NaCl, 0.5 mM EDTA) supplemented with protease inhibitor cocktail (6 μg/ml chymostatin, 0.5 μg/ml leupeptin, 10 μg/ml antipain, 2 μg/ml aprotinin, 0.7 μg/ml pepstatin A and 10 μg/ml 4-amidinophenylmethanesulfonyl fluoride hydrochloride; Sigma-Aldrich) and phosphatase inhibitor cocktails 2 and 3 (P5726 and P0044, Sigma-Aldrich). Protein concentration was measured with BCA Protein Assay Kit (Thermo Fisher Scientific, Cat no 23225). Subsequently, 15-25 μg of total protein per sample were resolved on 8%-14% SDS-PAGE and transferred onto nitrocellulose membrane (Amersham Hybond, GE Healthcare Life Science, 10600002). Membranes were blocked in 5% milk in PBS followed by incubation with specific primary and secondary antibodies. For signal detection, Clarity Western ECL Substrate (BioRad, 170-5061) and ChemiDoc imaging system (Bio-Rad) were applied. Densitometric analysis of western blotting bands was performed using Image Lab 6.0.1 software (Bio-Rad). The raw data were normalized to vinculin band intensities and presented as fold levels to the average of siCtrl#1 and siCtrl#2.

### Cell fractionation

Cellular fractionation was performed as described elsewhere (Suzuki *et al*, 2010). Briefly, cells growing on P100 dish were washed with ice-cold PBS, scraped and collected in 1.5 ml micro-centrifuge tube. After centrifugation (10 sec, 1.7×10^3^ g) pellet was resuspended in 900 μl of ice-cold 0.1% NP40 (IGEPAL® CA-630, I8896, Sigma Aldrich) in PBS and 300 μl of the lysate (whole cell lysate fraction, W) was transferred to a separate tube. The remaining material was centrifuged (10 sec, 1.2×10^4^ g), and the pellet was resuspended in 1 ml of ice-cold 0.1% NP40 in PBS and centrifuged (10 sec, 1.2×10^4^ g). Pellet (~20 μl) was resuspended in 180 μl of 1 × Laemmli sample buffer (nuclear fraction, N). Lysates were sonicated and boiled for 1 min 95° C.

### Quantitative real-time PCR (qRT-PCR)

Total RNA was isolated from cells with High Pure Isolation Kit (Roche, 11828665001) according to the manufacturer’s instruction. For cDNA preparation, 500 ng of total RNA, random nonamers (Sigma-Aldrich, R7647), oligo(dT)23 (Sigma-Aldrich, O4387) and M-MLV reverse transcriptase (Sigma-Aldrich, M1302) were used. Primers were designed using the NCBI Primer designing tool and custom-synthesized by Sigma-Aldrich. The sequences of primers were listed in **Supplementary Table**. For 2-3 technical repeats for each experimental conditions, cDNA sample amplification was performed with the KAPA SYBR FAST qPCR Kit (KapaBiosystems, KK4618) using the 7900HT Fast Real-Time PCR thermocycler (Applied Biosystems). Obtained data were normalized according to the expression level of the GAPDH (glyceraldehyde 3-phosphate dehydrogenase) housekeeping gene. Results are presented as fold change compared to siCtrl#1.

### Transcriptome analysis by RNA sequencing (RNA-Seq)

Transcriptome analysis of RKO cells was performed as described elsewhere (Kolmus **et al**, 2021). Briefly, cell pellet was collected 72 h post transfection with siRNAs. To generate sequencing library, Ion AmpliSeq Transcriptome Human Gene Expression Panel (ThermoFisher Scientific) was used. Sequencing was performed with Ion PI Hi-Q Sequencing 200 Kit (ThermoFisher Scientific) using Ion Proton instrument. Alignment of reads to the hg19 AmpliSeq Transcriptome ERCC v1 was performed with Torrent Mapping Alignment Program (version 5.0.4, ThermoFisher Scientific), followed by transcript quantification with HTseq-count (version 0.6.0). Differential gene expression analysis was performed for genes with more than 100 counts across conditions using the R package DESeq2 (version 1.18.1; (Love *et al*, 2014)). Non-protein coding genes were excluded from the analysis. The genes, which overlapped for on-target siRNAs, with expression levels normalized to those in siCtrl#1-transfected cells, were applied to GO analysis of biological processes using clusterProfiler (version 3.6.0; (Yu *et al*, 2012)) and corrected for multiple testing using the Benjamini-Hochberg method. Only genes with adjusted p-value < 0.05 were considered as significant. Obtained counts were transformed using the Transcript Per Million (TPM) normalization method and converted to obtain Z-scores. To reduce the redundancy of terms, 0.6 cutoff was applied. Heatmaps of differentially expressed genes were generated using ComplexHeatmap (version 1.17.1; (Gu *et al*, 2016)). The above-mentioned calculations and visualizations were performed in R version 3.4.4 (https://www.R-project.org).

### Statistical analysis

Data are shown as mean +/− standard error of mean (SEM) from at least three independent biological experiments. Statistical analysis was performed using the Prism 8.4.3 (GraphPad Software) using unpaired two-tailed Student t test (for qRT-PCR analysis, western blotting densitometry and % of cells with TFEB or TFE3 in the nucleus from confocal microcopy analysis) or paired two-tailed Student t-test (for quantified parameters from confocal microcopy analysis representing fluorescence intensity expressed in arbitrary units (a.u.) and mean structures area). The significance of mean comparison is annotated as follows: ns, non-significant (P≥0.05) or indicated with exact p-value, *P<0.05, **P<0.01, ***P<0.001, ****P<0.0001. Results were considered significant when P<0.05.

## Acknowledgements

We thank Kamil Jastrzębski, Agata Poświata, Lidia Wolińska-Nizioł and Daria Zdżalik-Bielecka for critical reading of the manuscript.

## Author contributions

The research was conceived by MW, JC and MMiaczynska. Funding was acquired by MMiaczynska. Experiments were designed by MW and JC, and performed mostly by MW with support from JC, ES and NBH. Transcriptomic analysis and visualization was performed by KK, KG, MD, AP and MMikula. The manuscript was written by MW and JC, and edited by MMiaczynska. JC and MMiaczynska supervised the work. All authors approved the manuscript.

## Data Availability

The RNA sequencing data have been deposited to Gene Expression Omnibus (GEO) under the accession number: GSE178665 (https://www.ncbi.nlm.nih.gov/geo/query/acc.cgi?acc=GSE178665).

---

**Supplementary Fig. 1.**
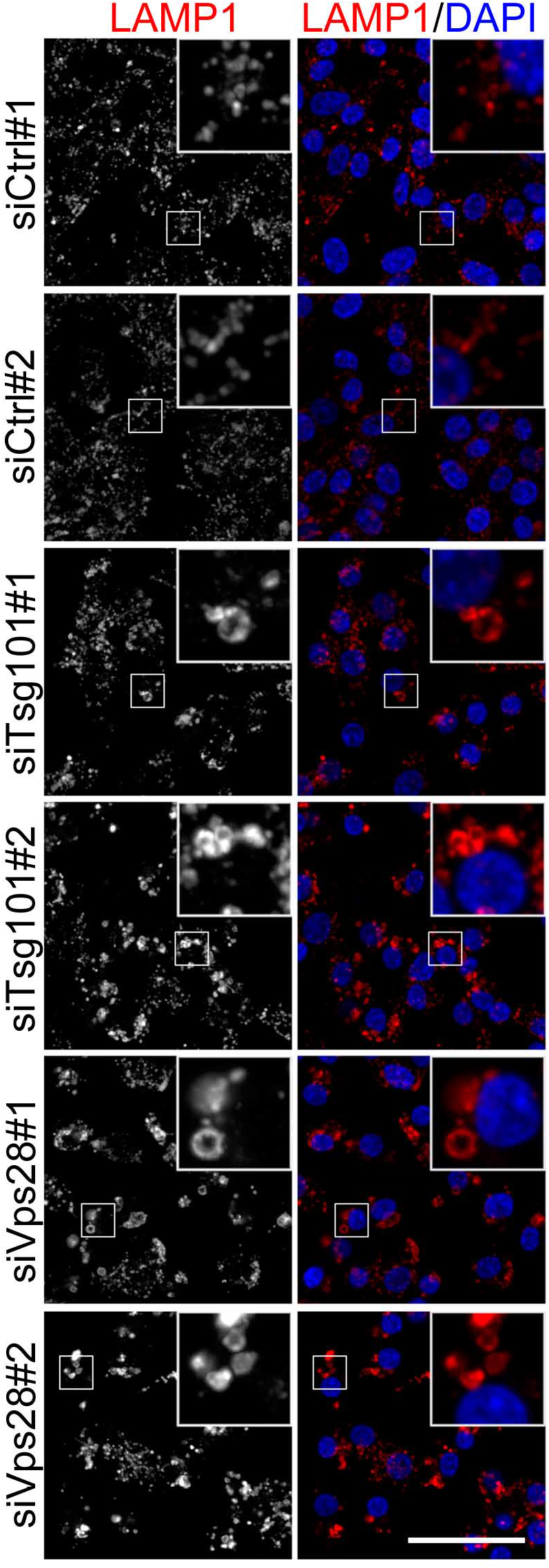
ESCRT-I depletion causes enlargement of LAMP1-positive structures in RKO cells. Representative maximum intensity projection confocal images of fixed RKO cells showing the effects of Tsg101 or Vps28 depletion (using two single siRNAs for each ESCRT-I subunit, #1 or #2) on the intracellular distribution of LAMP1 (red) as compared to control conditions (non-targeting siRNAs siCtrl#1 or #2). Cell nuclei marked with DAPI stain (blue). Scale bar, 50 μm.

**Supplementary Fig. 2.**
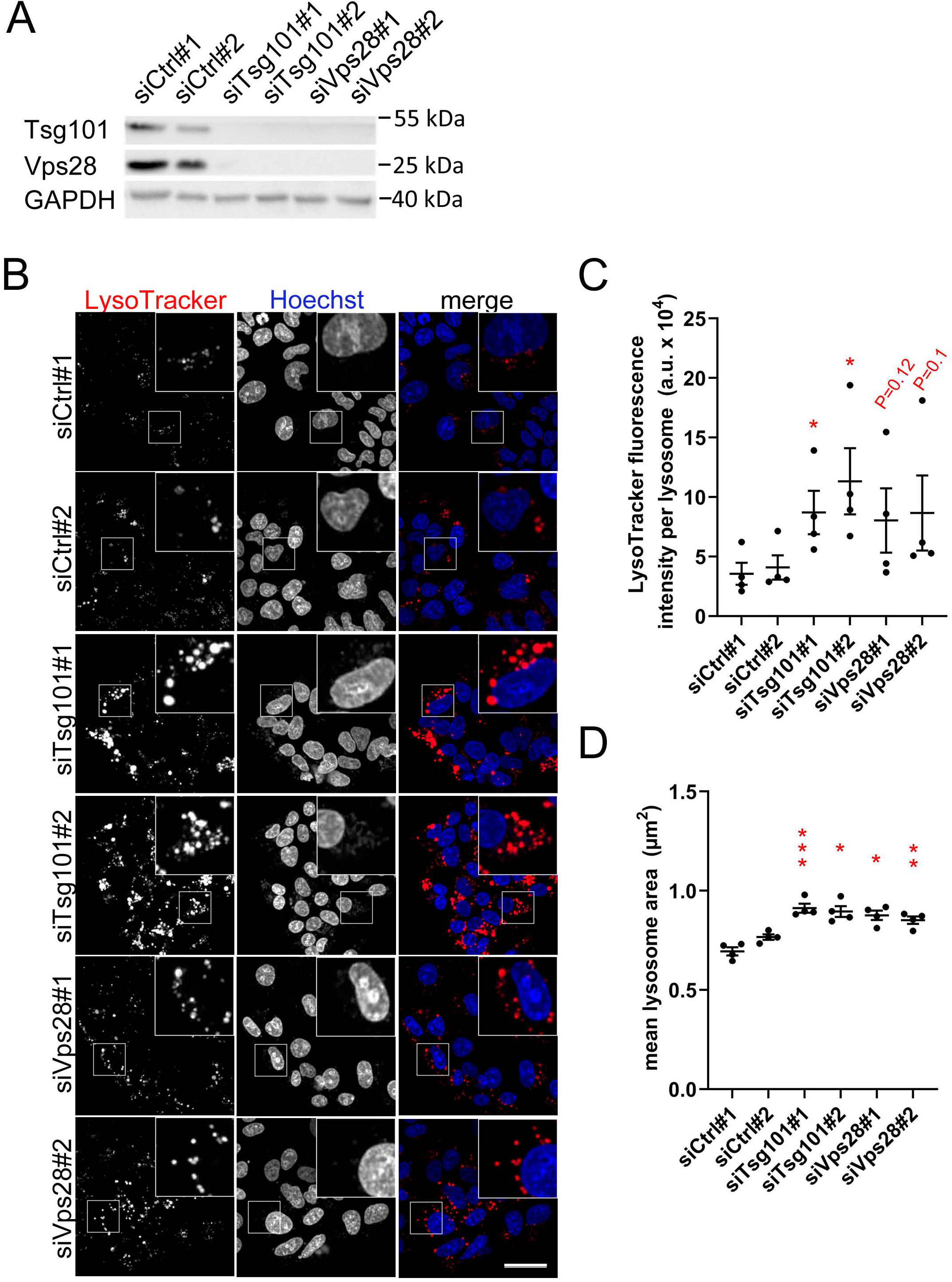
ESCRT-I dysfunction causes enlargement of lysosomes in DLD-1 cells. **(A)** Western blots showing the depletion efficiencies of ESCRT-I subunits, Tsg101 or Vps28 (using single siRNAs for each subunit, #1 or #2) as compared to control conditions (non-targeting siRNAs Ctrl#1 or #2). GAPDH was used as a loading control. **(B)** Maximum intensity projection confocal images of live cells, showing the intracellular distribution of lysosomes marked with LysoTracker dye (red) in control or ESCRT-I-depleted cells. Cell nuclei marked with Hoechst stain (blue). Scale bar, 50 μm. **(C-D)** Dot plots showing average fluorescence intensities of LysoTracker expressed in arbitrary units (a.u., **C**) and average area (**D**) of detected lysosomal structures in control or ESCRT-depleted cells, calculated based on live cell microscopy images (shown in B). Values derived from independent experiments (dots) and their means (n=4 +/− SEM) are presented. Statistical significance tested by comparison to averaged values measured for siCtrl#1 and #2. *P <0.05, **P<0.01, ***P<0.001.

**Supplementary Fig. 3.**
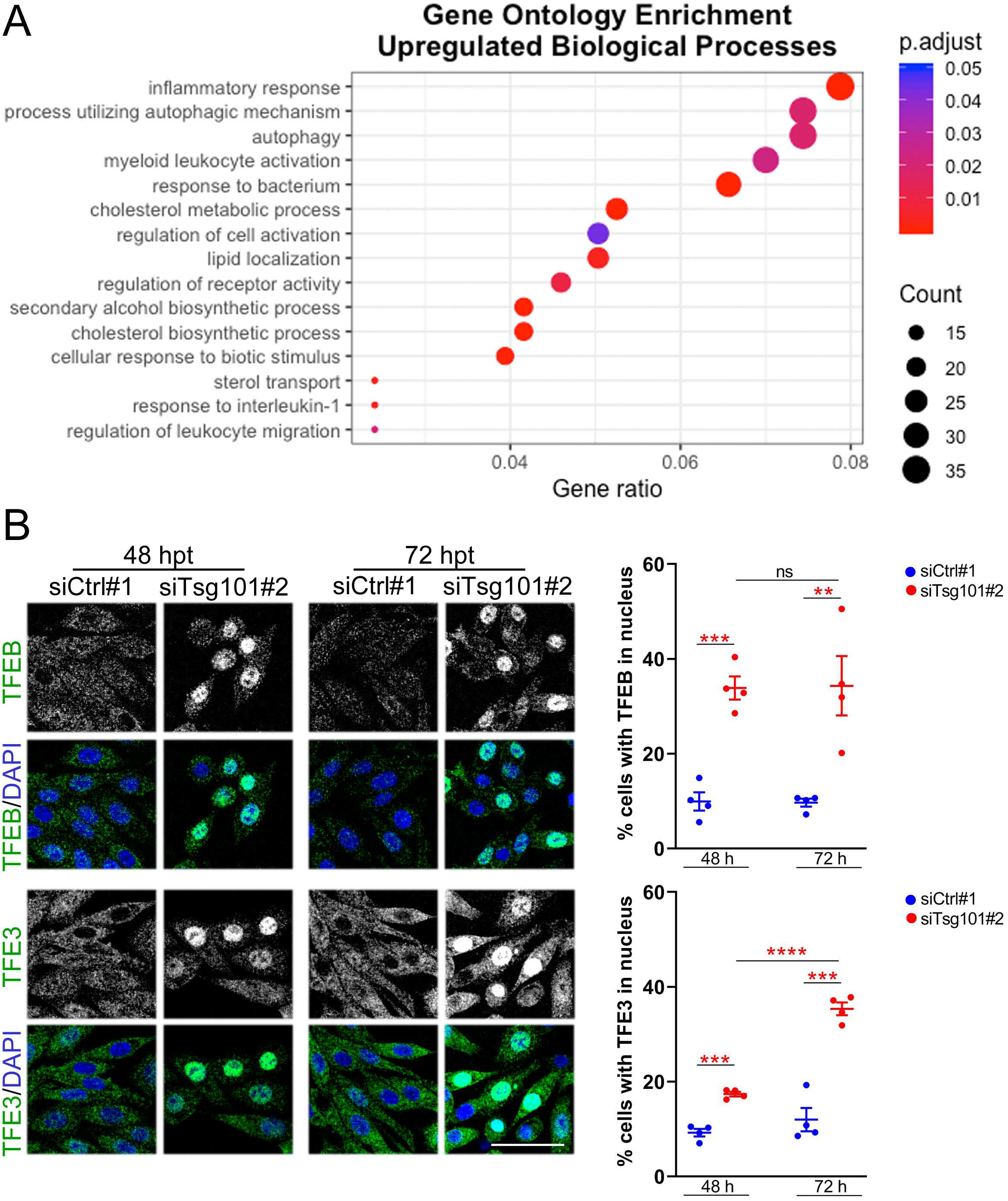
ESCRT-I deficiency induces signaling pathways indicative of lysosomal dysfunction. **(A)** Gene ontology (GO) analysis of top biological processes identified by annotation of genes with upregulated expression (≥ 1.5-fold; adjusted P-value < 0.05), detected by RNA-seq in RKO cells depleted of Tsg101 or Vps28 (siTsg101#1 or #2, siVps28#1 or #2) as compared to control cells (treated with non-targeting siRNAs, Ctrl#1 or #2). RNA-Seq data analysis was performed based on 3 independent experiments. **(B)** Representative maximum intensity projection confocal images of fixed RKO cells 48 hpt or 72 hpt (left panels) showing intracellular distribution of TFEB or TFE3 (green) in ESCRT-I-depleted (siTsg101#2) or control (Ctrl#1, non-targeting siRNA) RKO cells. Dot plots (right panels) showing percentage of cells with nuclear TFEB or TFE3 localization. Values derived from independent experiments (dots) and their means (n=4 +/− SEM) are presented. Statistical significance tested by comparison to siCtrl#1 and/or siTsg101#2 conditions. ns—non-significant (P≥0.05), **P<0.01, ***P<0.001, ****P<0.0001.

**Supplementary Fig. 4.**
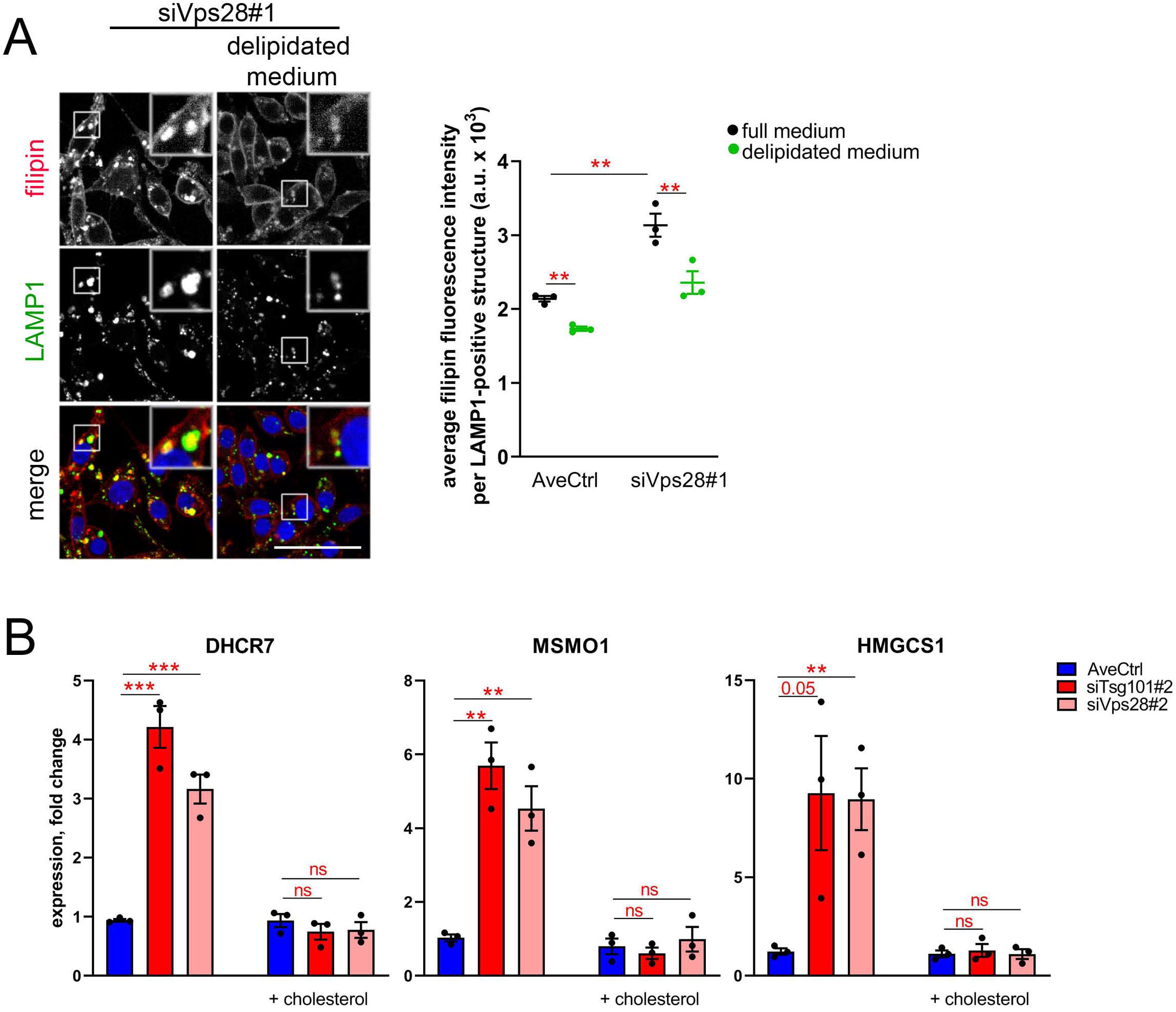
ESCRT-I deficiency upregulates expression of cholesterol biosynthesis genes due to impaired cholesterol efflux from lysosomes. **(A)** Representative maximum intensity projection confocal images of fixed RKO cells 48 hpt (left panel) showing the intracellular distribution of free cholesterol marked with filipin dye (red) with respect to LAMP1 (green) expressed in arbitrary units (a.u.) in Vps28-depleted cells and/or upon 40 h culture in a delipidated medium. Cell nuclei marked with DRAQ7 dye (blue). Scale bar, 50 μm. Dot plot (right panel) showing the average filipin fluorescence intensity per LAMP1-positive structure as compared to control cells, whose representative pictures are shown in **Fig. 4B**. Values derived from independent experiments (dots) and their means (n=3 +/− SEM) are presented. Statistical significance tested by comparison to averaged values measured for control cells (AveCtrl) and/or siVps28#1 conditions. Values for control cells are the same as in **Fig. 4B**.**P<0.01. **(B)** qPCR results showing the expression of cholesterol biosynthesis genes in ESCRT-I-depleted (siTsg101#2 or siVps28#2) or control (siCtrl#1,#2 non-targeting siRNAs) RKO cells 72 hpt and/or upon 48 h supplementation with water-soluble cholesterol (20 μM). Data are presented as fold changes with respect to siCtrl#1. Values derived from independent experiments (dots) and their means (n=3 +/− SEM) are presented. Statistical significance tested by comparison to averaged values measured for control cells (AveCtrl). ns—non-significant (P≥0.05), **P<0.01, ***P<0.001.

**Supplementary Table.**
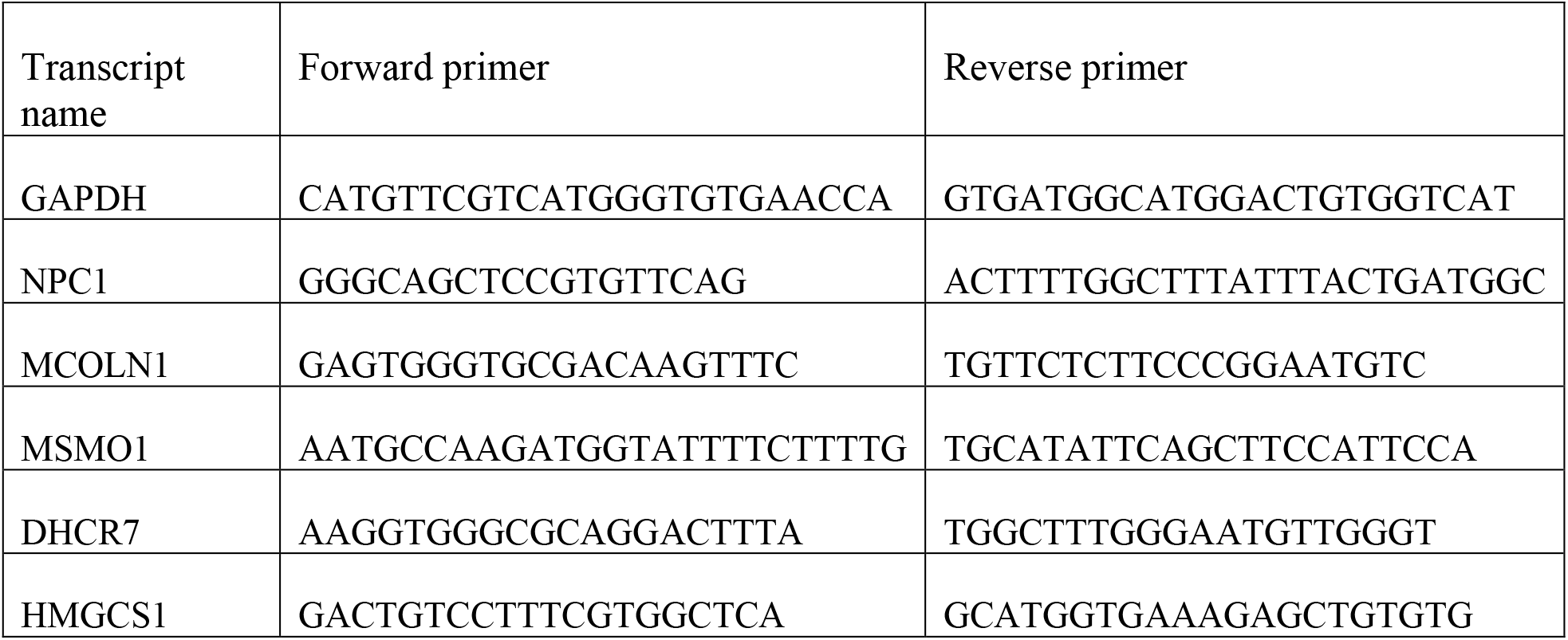
Sequences of primers used for qRT-PCR

